# Revised estimates for the number of human and bacteria cells in the body

**DOI:** 10.1101/036103

**Authors:** Ron Sender, Shai Fuchs, Ron Milo

**Affiliations:** Department of Plant and Environmental Sciences, Weizmann institute of science, Rehovot, Israel.; Department of Molecular Genetics, Weizmann institute of science, Rehovot, Israel.

## Abstract

We critically revisit the “common knowledge” that bacteria outnumber human cells by a ratio of at least 10:1 in the human body. We found the total number of bacteria in the “reference man” to be 3.9·10^13^, with an uncertainty (SEM) of 25%, and a variation over the population (CV) of 52%. For human cells we identify the dominant role of the hematopoietic lineage to the total count of body cells (≈90%), and revise past estimates to reach a total of 3.0·10^13^ human cells in the 70 kg “reference man” with 2% uncertainty and 14% CV. Our analysis updates the widely-cited 10:1 ratio, showing that the number of bacteria in our bodies is actually of the same order as the number of human cells. Indeed, the numbers are similar enough that each defecation event may flip the ratio to favor human cells over bacteria.

## Introduction

Study of the human microbiome has emerged as an area of utmost interest. The last two decades have produced an avalanche of studies revealing the impact that the microbiota have on the physiology and metabolism of multicellular organisms, with implications for health and disease. One of the most fundamental and commonly cited figures in this growing field is the estimate that bacteria residing in the human body outnumber human cells by a factor of 10 or more (Bäckhed et al., 2005; Gill et al., 2006; Round and Mazmanian, 2010; Turnbaugh et al., 2007). This striking statement often serves as an entry point to the field. After all, if a human being is a cell population composed of at least 90% bacteria, it is only natural to expect a major role for them in human physiology.

Both the numerator (number of microbial cells) and the denominator (human cells) of this 10:1 ratio are based on crude assessments. Most sources cite the number of human cells as 10^13^ or 10^14^, and a recent study reported 3.7·10^13^ human cells in a “reference” human (Bianconi et al., 2013). Estimates for the number of microbial cells in the body (which we operationally refer to as bacteria as they overwhelmingly outnumber eukaryotes and archaea in the human microbiome by 2 to 3 orders of magnitude) are usually 10^14^–10^15^ (Berg, 1996; Savage, 1977). We performed a thorough review of the literature and found a long chain of citations originating from one “back of the envelope” estimate. This estimate, though illuminating, was never meant to serve as the cornerstone of an entire field.

The aim of this study is to critically revisit the 10:1 estimate that has been so thoroughly repeated as to achieve the status of an established common knowledge fact. Recently this ratio was criticized in a letter to the journal Microbe (Rosner, 2014) An alternative estimate that will give concrete values and estimate the uncertainties range is needed. Here, we review the methodologies employed hitherto for cell count and perform a revised estimate. Doing so we repeat and reflect on the assumptions in previous back of the envelope calculations, also known as Fermi problems. We find such estimates as effective sanity checks and a way to improve our quantitative understanding in the biological context.

A major part of the available literature used in the derivation of human cell numbers was based on cohorts of exclusively or mostly men. As mentioned in the discussion, quantitative differences will apply for women due to changes in characteristic body mass, blood volume and the vaginal microbiota, but these are on the order of the reported uncertainties, and thus the paper conclusions are relevant for the general human population. The standard reference man we analyze is defined in the literature (Snyder et al., 1975) as: “Reference Man is defined as being between 20–30 years of age, weighing 70 kg, is 170 cm in height”. Our analysis revisits the estimates for the number of microbial cells, human cells and their ratio in the body of such a standard man.

We begin by revisiting the number of bacteria through surveying earlier sources, comparing counts in different body organs and finally focusing on the content of the colon. We then estimate the total number of human cells in a body comparing calculations using a “representative” cell size to aggregation by cell type, and contrasting the cell number distribution to the mass distribution. In closing we revisit the ratio of bacterial to human cells and evaluate the effect of age and gender.

## Results

### Origin of prevalent claims in the literature on the number of bacterial cells in humans

Bacteria are found in many parts of the human body primarily on the external and internal surfaces, including the gastrointestinal tracts, skin, saliva, oral mucosa and conjunctiva. The vast majority of commensal bacteria reside in the colon, with previous estimates of about 10^14^ bacteria (Savage, 1977), followed by the skin, which is estimated to harbor ~10^12^ bacteria (Berg, 1996). Because the number of bacteria in the gut dominates the total number of bacteria in the body, as discussed below in conjunction with Table 1, the current section reviews previous work to explain how the size of the bacterial population in the human gut has been estimated in the past. Almost all recent papers in the field of gut microbiota directly or indirectly rely on a single paper (Savage, 1977) regarding the overall number of bacteria in the gut. This is shown in Figure 1 as a citation lineage for a few representative cases. Interestingly, those who read the original paper (Savage, 1977) find that it actually cites another paper for his estimate (Luckey, 1972). This older paper performed an order-of-magnitude estimate by assuming 10^11^ bacteria per gram (which has some support in the literature) and 1 liter (or about 1 kg) of alimentary tract capacity. The estimate, performed by Luckey in 1972 is an illuminating example of back of the envelope estimate, which was elegantly performed, yet was probably never meant to be widely quoted decades later. More recently, a report from the NIH stated a value of 1–3% of the body mass is composed of bacteria (Macdougall, 2012). This value, quoted on Wikipedia (Wikipedia, 2015) among many other online resources, coupled with a rule of thumb for bacterial cell volume of 1 μm^3^ leads to an estimate of 10^15^ bacteria in the human body, which led to claims of 100:1 bacterial to human cells. After showing the dominance of gut bacteria we will revisit estimates of the number of bacteria in the human body.

**Figure 1:**
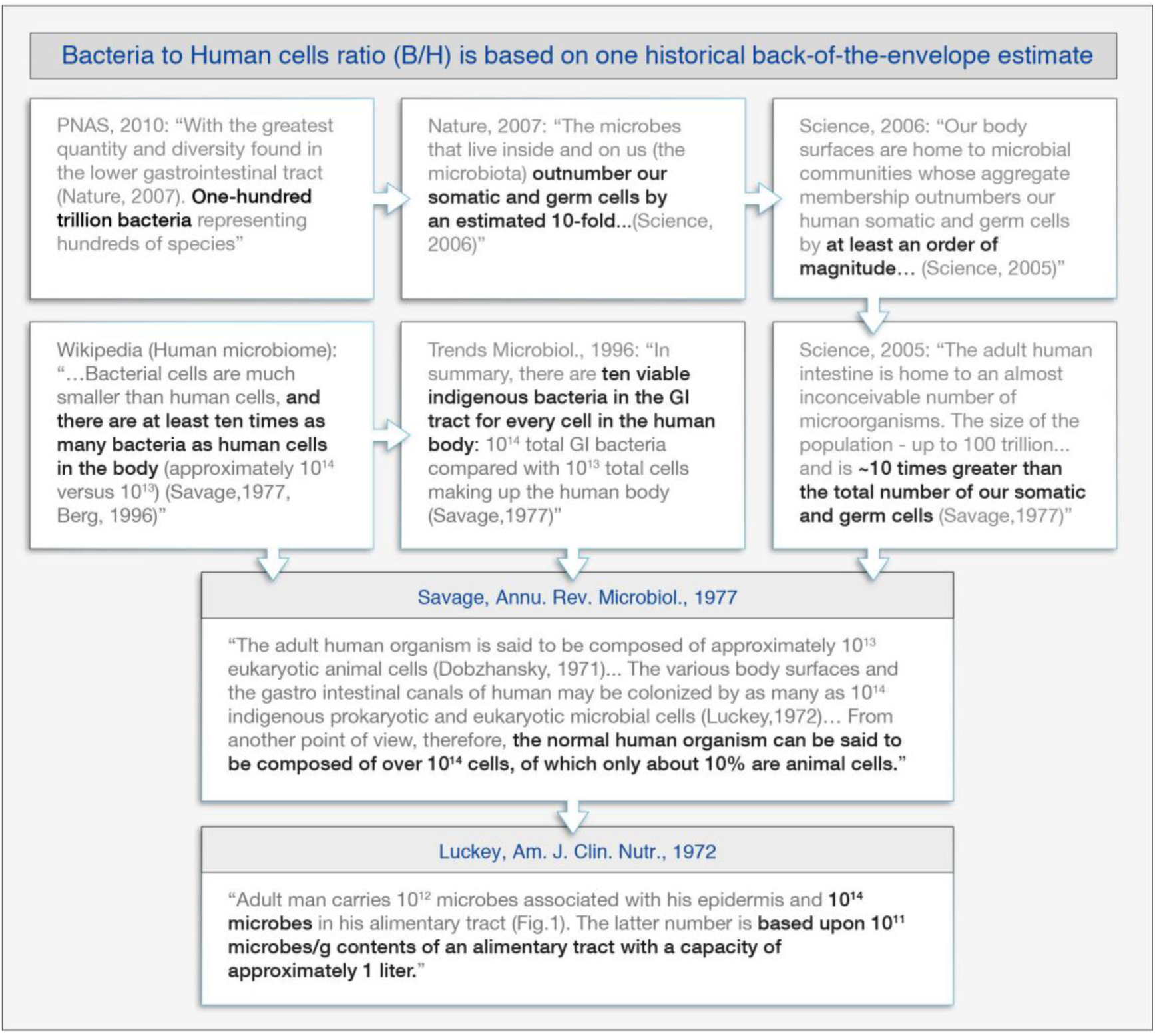
A non-exhaustive lineage tree of quotations showing the origins of the often-quoted sources for the number of bacteria in the human gut. The 1977 review by Savage is referenced over 1000 times in the literature often in the context of the estimate for the vast overabundance of bacteria over human cells. Brief quotes from the original papers are shown. Arrows point to the reference used. The numerical statements are in bold.

### Distribution of bacteria in different human organs

Table 1 shows typical order-of-magnitude estimates for the number of bacteria that reside in different organs in the human body. The estimates are based on multiplying measured concentrations of bacteria by the volume of each organ (Berg, 1996; Tannock, 1995). Values are rounded up to give an order of magnitude upper bound. For the skin we used bacterial areal density and total skin surface to reach an upper bound.

**Table 1:**
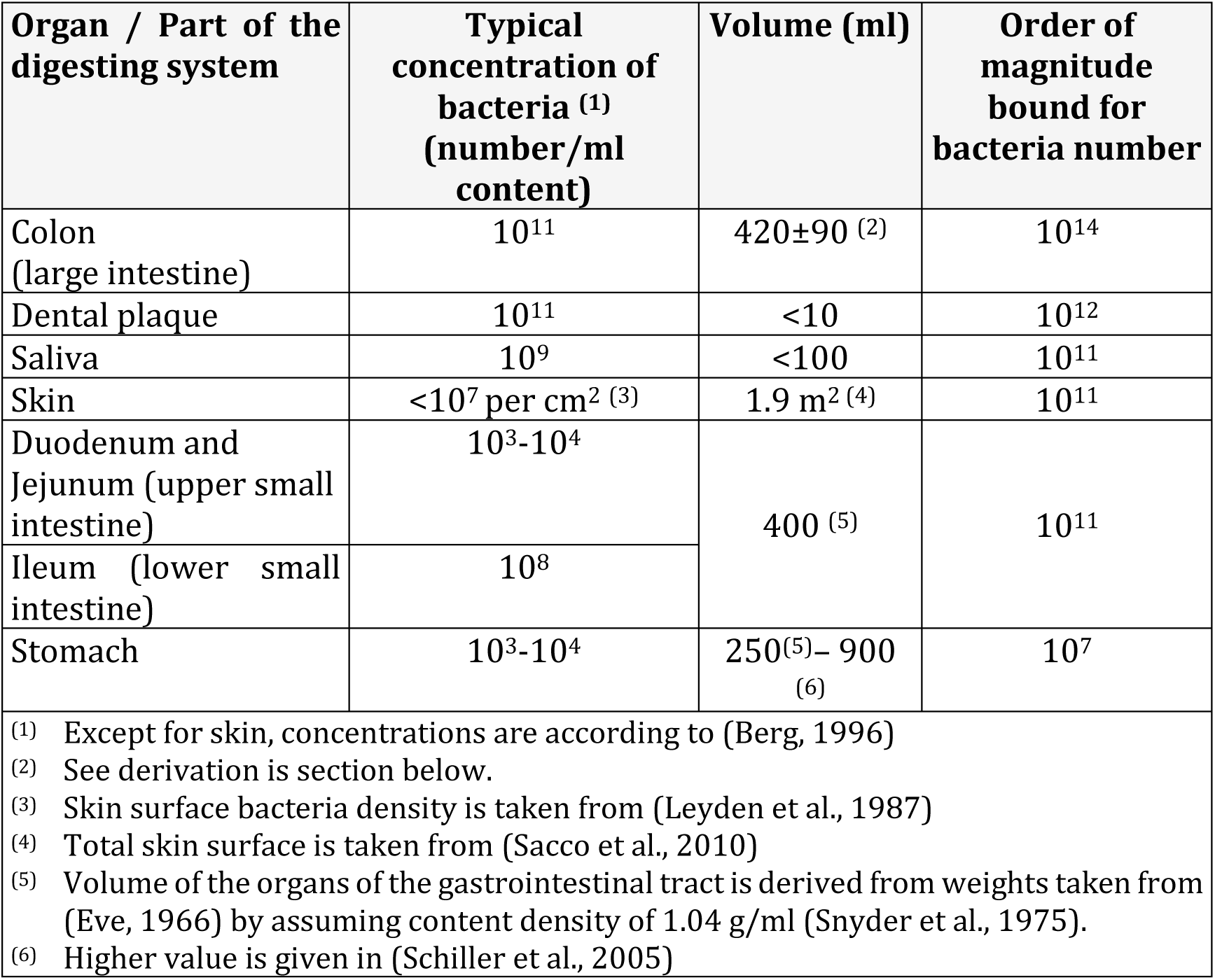
Bounds for bacteria number in different organs, derived from bacterial concentrations and volume.

| Organ / Part of the digesting system | Typical concentration of bacteria <sup>(1)</sup> (number/ml content) | Volume (ml) | Order of magnitude bound for bacteria number |
| --- | --- | --- | --- |
| Colon (large intestine) | $10^{11}$ | $420 \pm 90$ <sup>(2)</sup> | $10^{14}$ |
| Dental plaque | $10^{11}$ | $<10$ | $10^{12}$ |
| Saliva | $10^9$ | $<100$ | $10^{11}$ |
| Skin | $<10^7$ per $\text{cm}^2$ <sup>(3)</sup> | $1.9 \text{ m}^2$ <sup>(4)</sup> | $10^{11}$ |
| Duodenum and Jejunum (upper small intestine) | $10^3$-$10^4$ | $400$ <sup>(5)</sup> | $10^{11}$ |
| Ileum (lower small intestine) | $10^8$ | | |
| Stomach | $10^3$-$10^4$ | $250^{(5)} - 900^{(6)}$ | $10^7$ |
| <sup>(1)</sup> Except for skin, concentrations are according to (Berg, 1996)<br><sup>(2)</sup> See derivation in section below.<br><sup>(3)</sup> Skin surface bacteria density is taken from (Leyden et al., 1987)<br><sup>(4)</sup> Total skin surface is taken from (Sacco et al., 2010)<br><sup>(5)</sup> Volume of the organs of the gastrointestinal tract is derived from weights taken from (Eve, 1966) by assuming content density of 1.04 g/ml (Snyder et al., 1975).<br><sup>(6)</sup> Higher value is given in (Schiller et al., 2005) |  |  |  |

Although the bacterial concentrations in the saliva and dental plaque are high, because of their small volume the overall numbers of bacteria in the mouth represents less than 1% of the colon bacterial content. The concentration of bacteria in the stomach and the upper 2/3 of the small intestine (duodenum and jejunum) is only 10^3^–10^4^ bacteria/ml, owing to the relative low pH of the stomach and the fast flow of the content through the stomach and the small intestine (Tannock, 1995). Table 1 reveals that the bacterial content of the colon exceeds all other organs by at least two orders of magnitude. Importantly, within the alimentary tract the colon is the only substantial contributor to the total bacterial population, while the stomach and small intestine make negligible contributions.

### Revisiting the original back-of-the-envelope estimate for the number of bacteria in the colon

The often cited value of ~10^14^ bacteria in the body has its roots in the 1970’s (Holdeman et al., 1976; Luckey, 1972; Moore and Holdeman, 1974). Figure 1 shows a few representative cases of how current citations converge to one original source. The first estimate of the number of bacteria in the human colon (Luckey, 1972), which we identify as the primary reference across the literature, was made by taking the volume of the alimentary tract, assumed as 1 liter, and multiplying it by the number density of bacteria, assumed to be 10^11^ bacteria per gram of wet content. Such estimates are often very illuminating, yet it is useful to revisit them as more empirical data accumulates. This pioneering estimate of 10^14^ bacteria in the colon, is based on a constant bacterial density in 1 Liter of alimentary tract volume (converting from volume to mass via a density of 1 g/ml). However, the number of bacteria in the alimentary tract proximal to the colon is negligible in comparison to the colon content (Table 1). Thus the relevant volume for the 10^11^ bacteria/g density is only that of the colon.

The inner volume of the colon of the reference adult man was estimated as 340 ml (355 g at density of 1.04 g/ml (Snyder et al., 1975)) based on various methods including flow measurements, barium meal X-ray measurements and post-mortem examination (Eve, 1966). A recent study The gives data about the volume of undisturbed colon that was gathered by MRI scans. The height-standardized regional colonic inner volume index for men is given to be 97±24 ml/m^3^ (±SD) dividing the colonic volume by the cube of the height in meters. Taking a height of 1.70 m for the reference man (Snyder et al., 1975), we arrive at a colon volume of 477±118 ml (SD). We can sanity-check this volume estimate by looking at the volume of stool that flows through the colon. An adult human is reported to produce 147 grams of wet stool per day (SEM 16%) (Hammer et al., 1997; Stephen et al., 1987; Wyman et al., 1978). The normal colonic transit time is about 34 hours (SEM 10%) (Southwell et al., 2009). We thus get a volume estimate of 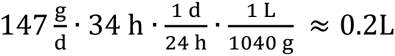 (SEM 20%) which is somewhat lower than but consistent with the values above given the uncertainties and very crude estimate. To summarize, the volume of colon content has been measured by two studies and independently validated by considering fecal transit dynamics. Values for a reference adult man averaged 409 ml (SEM 16%, CV 25%), which will be used in calculations below.

### Concentration of bacteria in the colon

The most widely used approach for measuring the bacterial cell density in the colon is by examining bacteria content in stool samples. This assumes that stool samples give adequate representation of colon content. We return to this assumption in the discussion. The first such experiments date back to the 1960’s and 1970’s (Holdeman et al., 1976; Houte and Gibbons, 1966; Moore and Holdeman, 1974; Stephen and Cummings, 1980). Researchers took stool samples from several patients and examined the bacteria number density in them. To do so, they used direct microscopic clump counts from diluted samples. Later experiments (Franks et al., 1998; Harmsen et al., 2002; Tannock et al., 2000; Thiel and Blaut, 2005) used DAPI staining and FISH (fluorescent in situ hybridization). Values are usually reported as bacteria per gram of dry stool. However, for our calculation we are interested in the bacteria content for the wet rather than dry content of the colon. To move from 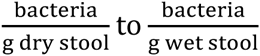 we use the fraction of dry matter as reported in each article. Table 2 reports the values we were able to find in the literature and translated them to a common basis.

**Table 2:** Values of bacteria density in stool as reported in several past articles. Mean bacteria number is calculated using the geometric mean to give robustness towards outlier values. Values quoted directly from the articles are written in bold, values derived by us are written in italic. Values reported with more than 2 significant digits are rounded to two significant digits as the uncertainty makes such over specification non sensible. ^#^Value for (Stephen and Cummings, 1980) derived from their table 1. *From derivation, assuming the averaged dry matter fraction of 26%. ± standard error of the mean

| Article | **bac. #/g**<br>**dry stool**<br>**(x10<sup>11</sup>)** | **dry matter**<br>**as % of**<br>**stool** | **bac. #/g**<br>**wet stool**<br>**(x10<sup>11</sup>)** | **CV(%)** |
| --- | --- | --- | --- | --- |
| (Houte and Gibbons, 1966) | - | - | **3.2** | 53% |
| (Moore and Holdeman, 1974) | **5** | **22%** | **1.1** | 78% |
| (Holdeman et al., 1976) | **4.1** | **31%** | *1.3* | 66% |
| (Stephen and Cummings, 1980) | **4** | - | *1.2<sup>#</sup>* | 25% |
| (Langendijk et al., 1995) | - | - | **2.7** | 26% |
| (Franks et al., 1998) | **2.9** | - | *0.76<sup>\*</sup>* | 39% |
| (Simmering and Kleessen, 1999) | **4.8** | - | *1.3<sup>\*</sup>* | 44% |
| (Tannock et al., 2000) | - | - | **0.95** | 40% |
| (Harmsen et al., 2002) | **2.1** | *30%* | **0.62** | 38% |
| (Zoetendal et al., 2002) | **2.9** | - | *0.76<sup>\*</sup>* | 24% |
| (Zhong et al., 2004) | **1.5** | **23%** | *0.35* | 73% |

| Article | bac. #/g<br>dry stool<br>($\times 10^{11}$) | dry matter<br>as % of<br>stool | bac. #/g<br>wet stool<br>($\times 10^{11}$) | CV(%) |
| --- | --- | --- | --- | --- |
| (Thiel and Blaut, 2005) | 3.5 | 25% | 0.87 | 53% |
| (He et al., 2008) | 1.5 | - | 0.38* | 43% |
| (Uyeno et al., 2008) | - | - | 0.44 | 34% |
| Mean | - | 26%±2% | 0.92±19% | 46% |

From the measurements collected in Table 2 we calculated the representative bacteria concentration in the colon by two methods, yielding very close values: the geometric mean is 0.92·10^11^ (SEM 19%) bacteria per gram wet stool, while median of the values is 0.91·10^11^ (bootstrapping SEM 19%, see methods). The Variation of the population, given by the average CV is 46%.

What fraction of colonic content is occupied by bacterial mass? What is the mean mass of a bacterium in the colon? The above measurements can be used to infer answers to those questions, provided two additional values: (1) the fraction of dried fecal mass that is dry bacteria and (2) the total water content in a bacterium. Overall dry mass fraction contributed by bacteria was directly measured to be 55% of fecal dry mass (Stephen and Cummings, 1980). The dry mass percentage of cell mass varies for different types of bacteria (Bratbak and Dundas, 1984; Robertson and Button, 1998) but can be assumed to be roughly equal to that of stool (29%), and thus the fraction of bacterial dry mass in dry feces is a good approximation to the fraction of bacterial mass in stool. Using the measured value of 4·10^11^ bacteria per gram dry stool (Stephen and Cummings, 1980), we evaluate the average mass of bacteria in the Stephen and Cummings samples to be 4.6·10” ^12^ g (SEM 35%, CV 47%). Interestingly, this value for the average bacterial cell mass is several times higher than is usually taken for a model bacterium such as *E. coli* (Chesbro et al., 1979; Kubitschek et al., 1986).

### Updated estimate for number of bacteria in the colon

We are now able to repeat the original calculation for the number of bacteria in the colon (Luckey, 1972). Given 0.9·10^11^ bacteria/g wet stool and 0.41 L of colon we find 3.9·10^13^ bacteria in the colon with an uncertainty of 24% and a variation of 52% over a population of standard weight males. Considering that the contribution to the total number of bacteria from other organs is at most 10^12^, we use 3.9·10^13^ as our estimate for the number of bacteria in the “reference man”.

### The number of human cells in a “standard” adult male

In the literature we find many statements for the number of cells in the human body ranging from 10^12^ to 10^14^ cells (Alberts et al., 2002; Cooper and Hausman, 2000; Goodsell, 2009; Griffiths, 2005; Lodish, 2000). A mass-based order-of-magnitude estimate for this number assumes a 10^2^ kg man, which is divided by the mass of a “representative” mammalian, cell 10^−12^–10^−11^ kg (assuming cell volumes of 1,000–10,000 μm^3^, respectively), thus arriving at 10^13^ - 10^14^ cells. For these kind of estimates, where cell mass is estimated to within an order of magnitude, factors contributing to less than 2-fold difference are neglected. Thus we use 100 kg as the mass of a reference man instead of 70 kg and we ignore the contribution of extracellular mass to the total mass. These simplifications are useful in making the estimate concise and transparent.

One study used a DNA-centered method (Baserga, 1985). The method begins by estimating the number of cells in the body of a 25g mouse by dividing the total amount of DNA (stated to be 20 mg) by the DNA content of one diploid mouse cell (6·10^−12^ g DNA per cell) to get ≈3·10^9^ cells in a 25 g mouse. Then extrapolate from mouse to human by using the ratio of masses to get ≈10^13^ cells in the human body. This method excludes cells that do not contain DNA, such as red blood cells and platelets.

An alternative approach that bypasses the need to think of a representative “average” cell systematically counts cells by type. A detailed analysis of this sort was recently published (Bianconi et al., 2013). The number of cells in the body by type or organ system was estimated. Since the aim was to systematically scrutinize all cellular components, the authors alternate between grouping by histologic type (e.g. glial cells) or by locus/organ where both parenchymal and stromal cells are accounted for (e.g. “bone marrow nucleated cells”). For each category the cell count was obtained by either a literature reference or by a calculation based on direct count in histological cross sections. Summing over a total of 56 cell categories (Bianconi et al., 2013) resulted in an overall estimate of 3.7·10^13^ human cells in the body (SD 0.8·10^13^, i.e. CV of 22%).

### Updated inventory of human cells in the body

In our effort to revisit the measurements cited we employed an approach that tries to combine the detailed, census approach with the benefits of a heuristic calculation used as a sanity check. We focus on the six cell types that were recently identified (Bianconi et al., 2013) to comprise 97% of the human cell count: red blood cells (accounting for 70%), glial cells (8%), endothelial cells (7%), dermal fibroblasts (5%), platelets (4%) and bone marrow cells (2%). The other 50 cell types account for the remaining 3%. In four cases (red blood cells, glial cells, endothelial cells, and dermal fibroblasts) we suggest a revised calculation as detailed in this section.

The largest contributor to the overall number of cells are red blood cells. Calculation of the number of red blood cells was made (see SI tab RBC#) by taking an average blood volume of 4.9L (SEM 1.6%, CV 9%) (Boer, 1984; Feldschuh and Enson, 1977; Nadler et al., 1962; Snyder et al., 1975) multiplied by a mean RBC count of 5.0·10^12^ cells/L (SEM 1.2%, CV 7%) (Ambayya et al., 2014; Dosoo et al., 2012; Nordin et al., 2004; Pekelharing et al., 2010; Volkmer and Heinemann, 2011; Wakeman et al., 2007). The latter could be verified by looking at your routine complete blood count, normal values range from 4.6–6.1·10^12^ cells/L for males and 4.2–5.4·10^12^ cells/L for females. This led to a total of 2.5·10^13^ red blood cells (SEM 2%, CV 12%). This is similar to the earlier report of 2.6·10^13^ cells (Bianconi et al., 2013).

The number of glial cells was previously reported as 3·10^12^ (Bianconi et al., 2013). This estimate is based on a 10:1 ratio between glial cells and neurons in the brain. This ratio of glia:neuron was held as a broadly accepted convention across the literature. However, a recent analysis (Azevedo et al., 2009) revisits this value and after analyzing the variation in between brain regions concludes that the ratio is close to 1:1. The study concludes that there are 8.5·10^10^ glial cells (CV 11%) in the brain and a similar number of neurons and so we use these values here.

The number of endothelial cells in the body was earlier estimated at 2.5·10^12^ cells (CV 40%), by taking the mean surface area of one endothelial cell (Bianconi et al., 2013) and dividing it into the total surface area of the blood vessels, based on a total capillary length of 8·10^9^ cm. We could not find a primary source for the total length of the capillary bed and thus decided to revisit this estimate. We used data regarding the percentage of the blood volume in each type of blood vessels (Leggett and Williamst, 1991). Using mean diameters for different blood vessels (Burton, 1954) we were able to derive the total length of each type of vessel (arteries, veins, capillaries etc.) and its corresponding surface area. Dividing by the mean surface area of one endothelial cell (Félétou, 2011) we derive a total estimate of 6·10^11^ cells.

The number of dermal fibroblast was previously estimated to be 1.85·10^12^ (Bianconi et al., 2013), based on multiplying the total surface area of the human body (SA=1.85 m^2^ (Herman, 2007)) by an areal density of dermal fibroblasts (Randolph and Simon, 1998). We wished to incorporate the dermal thickness (d) into the calculation. Dermal thickness was directly measured at 17 points throughout the body (Moore et al., 2003), with the mean of these measurements yielding 0.11±0.04 cm. The dermis is composed of two main layers: papillary dermis (about 10% of the dermis thickness) and reticular dermis (the other 90%) (McGrath et al., 2004). The fibroblast density is greater in the papillary dermis, with a reported areal density, σ_pap_. of 10^6^ cells/cm^2^ (with 100 μm thickness of papillary, giving 10^8^ cells/cm^3^) (Randolph and Simon, 1998). The fibroblast density in the middle of the dermis was reported to be about 3·10^6^ cells/cm^3^ (Miller et al., 2003) giving an areal density of σ_ret_. = 3·10^5^ cells/cm^2^. Combining these we find: 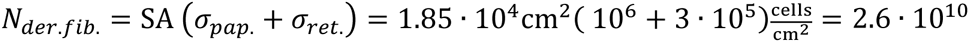 cells. Even after this 100-fold decrease in number, dermal fibroblasts are estimated to account for only ≈0.05% of the human cell count.

To summarize, of the previously estimated 9·10^12^ non-blood cells in a human body, our revised calculations yield only about 9·10^11^ cells. This revision reduces the number of non-blood cells to 3·10^12^, merely 10% of the total updated human cell count. The striking dominance of the hematopoietic lineage in cell count (90% of the total) is counterintuitive given the composition of the body by mass. This is the subject of the following analysis. In Figure 2 we summarize the revised results for the contribution of the different cell types to the total number of human cells. Categories contributing >0.4% are presented. All the other categories sum up to about 2% together.

**Figure 2:**
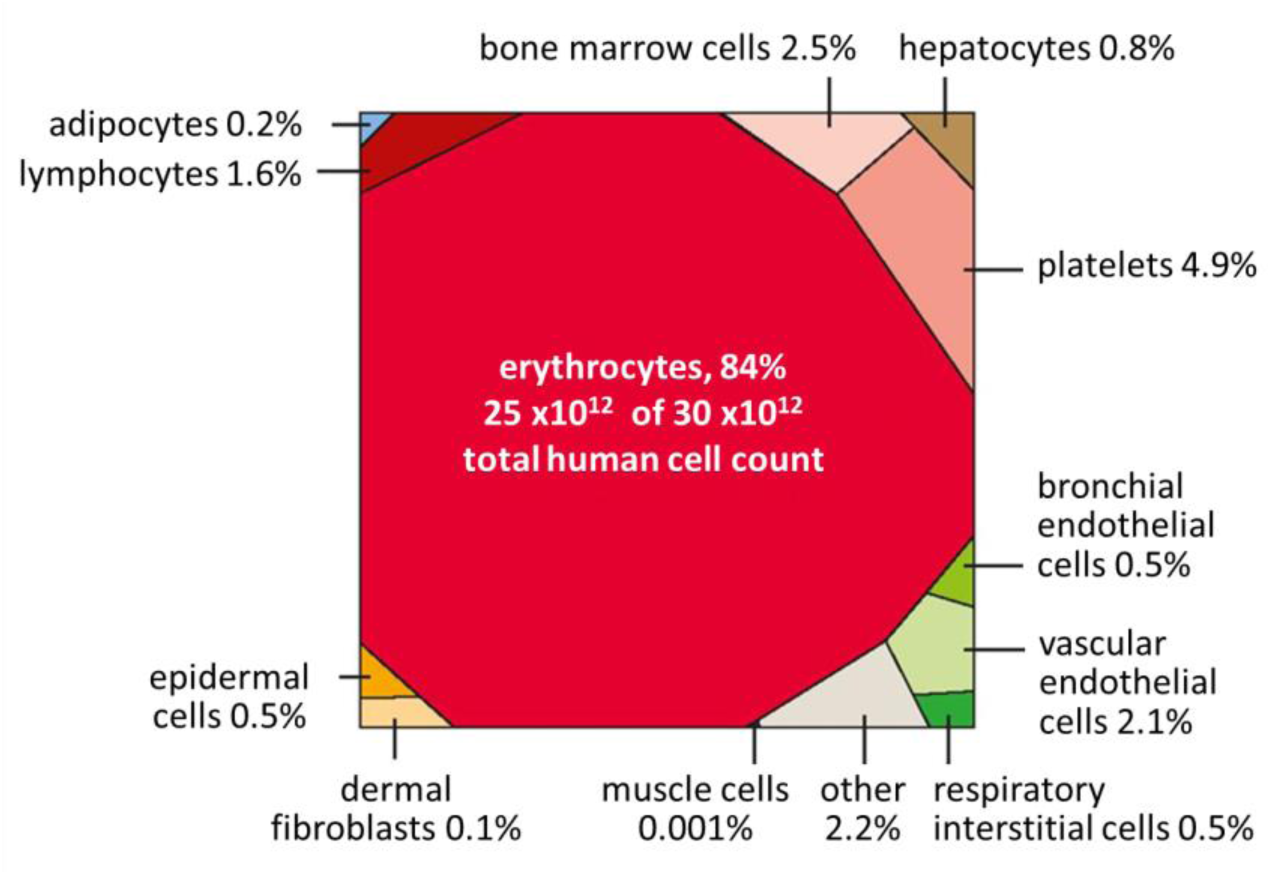
The distribution of number of cells by cell type. Note that muscles cells and fat cells, each comprising about 20 kg of tissue, have a small contribution to the total number of cells (0.2% or less) due to their large cell size.

The visualization in Figure 3 highlights that almost 90% of the cells are estimated to be enucleated cells (26·10^12^ cells), mostly red blood cells and platelets, circulating in the blood vessels.

**Figure 3:**
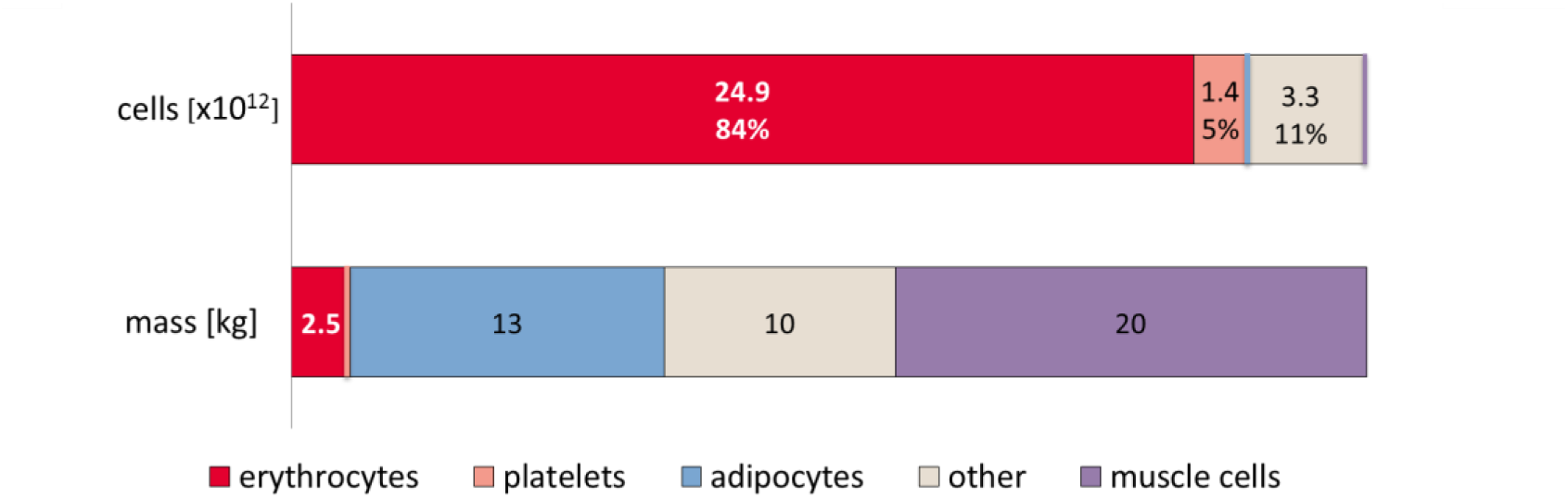
Distribution of cell number and mass for different cell types in the human body (for a 70 kg adult man). The upper bar displays the number of cells, while the lower bar displays the main cell types comprising the overall body mass.

### Mass centered approach as sanity check for cell count

Next we asked whether it is possible that a large bulk of cells was neglected due to significant underestimation. Have Bianconi et al. (Bianconi et al., 2013) accounted for the main bulk of cells in the body? One may naively assume that a way of approaching this question is through mass – does the cumulative mass of the cells counted fall within a reasonable range? To properly tackle that question we first need to state what the anticipated result is, i.e. total body cell mass. For a reference man mass of 70 kg, 25% is extracellular fluid (Shen et al., 2007), another 7% is extra-cellular solids (Shen et al., 2007), thus we need to account for ≈47 kg of cell mass (including fat).

A comprehensive systematic source for the composition of total cell mass (rather than total cell count) is the *Report of the Task Group on Reference Man* (Snyder et al., 1975) which gives values for the mass of the main tissues of the human body. This mass per tissue analysis includes both intra- and extra-cellular components. To distinguish between intra- and extra-cellular portions of each tissue we can leverage total body potassium measurements (Wang et al., 2004, 2005). The concentration of potassium in the intracellular and extracellular volumes of the body is known to be relatively constant (Wang et al., 2004). Given these constant values, Wang derived a formula connecting the potassium level of a tissue with its non-fat cell mass. The extracellular potassium concentration is only about 3% of the intracellular concentration and thus can be neglected to give the relation M_tissue_(kg)=0.0092 [K] (mmol). We used this relation to derive the cell mass in each of the main tissues from reported potassium concentrations (Snyder et al., 1975).

After this review and adjustment, what can we learn from connecting the tissue masses (Snyder et al., 1975) to detailed cell counts? Figure 3 compares the main tissues that contribute to the human body, in terms of cell number and masses.

A striking outcome of this juxtaposition is the evident discrepancy between contributors to total cell mass and to cell number. The cell count is dominated by red blood cells (84%), among the smallest cell types in the human body with a volume of about 100 μm^3^. In contrast, 75% of total cell mass is constituted by two cell types, fat cells (adipocytes) and muscle cells (myocytes), large cells (usually >10,000 μm^3^ by volume) that compose only a minute fraction (≈0.1%) of total cell number.

If there exists a bulk of cells large enough to alter the total cell count it should contain on the order of 10^12^ cells or more. However, these cells cannot have a total mass more than a few kg at the most, as the mass of the reference body is already almost fully accounted for. Therefore, any such cells would need to have rather small mass, with an upper bound of 1 kg/10^12^ cells <1,000 pg/cell. Thus we conclude that if indeed there is any underestimation or omission in the cell count, for it to have a sizeable effect on total count, it should be of small cells.

The hematopoietic lineage has generally small cells and is a good candidate for undercounting. Bianconi et al (Bianconi et al., 2013) accounted most of the cells of this lineage: red blood cells, white blood cells and platelets were counted in the blood and in the bone marrow, but there is a non-negligible fraction of the white blood cells that presented outside of these tissues. The total number of lymphocytes in a 70 kg man is estimated to be about 5·10^11^ (Trepel, 1974), most of them reside in the lymphatic system and in tissues across the body. Thus, while Bianconi et al. (Bianconi et al., 2013) have not considered lymphocytes outside the blood and bone marrow in their account, this underestimated population has only a marginal effect on the total cell count.

Figure 4 summarizes our revised cell count and tissue mass data for easy cross-reference. We conclude that, total count of cells in our body is divided between ≈3 10^12^ nucleated cells which account for ≈10% of the cells, while cells from the hematopoietic lineage comprise nearly 90% of total human cell count.

**Figure 4.**
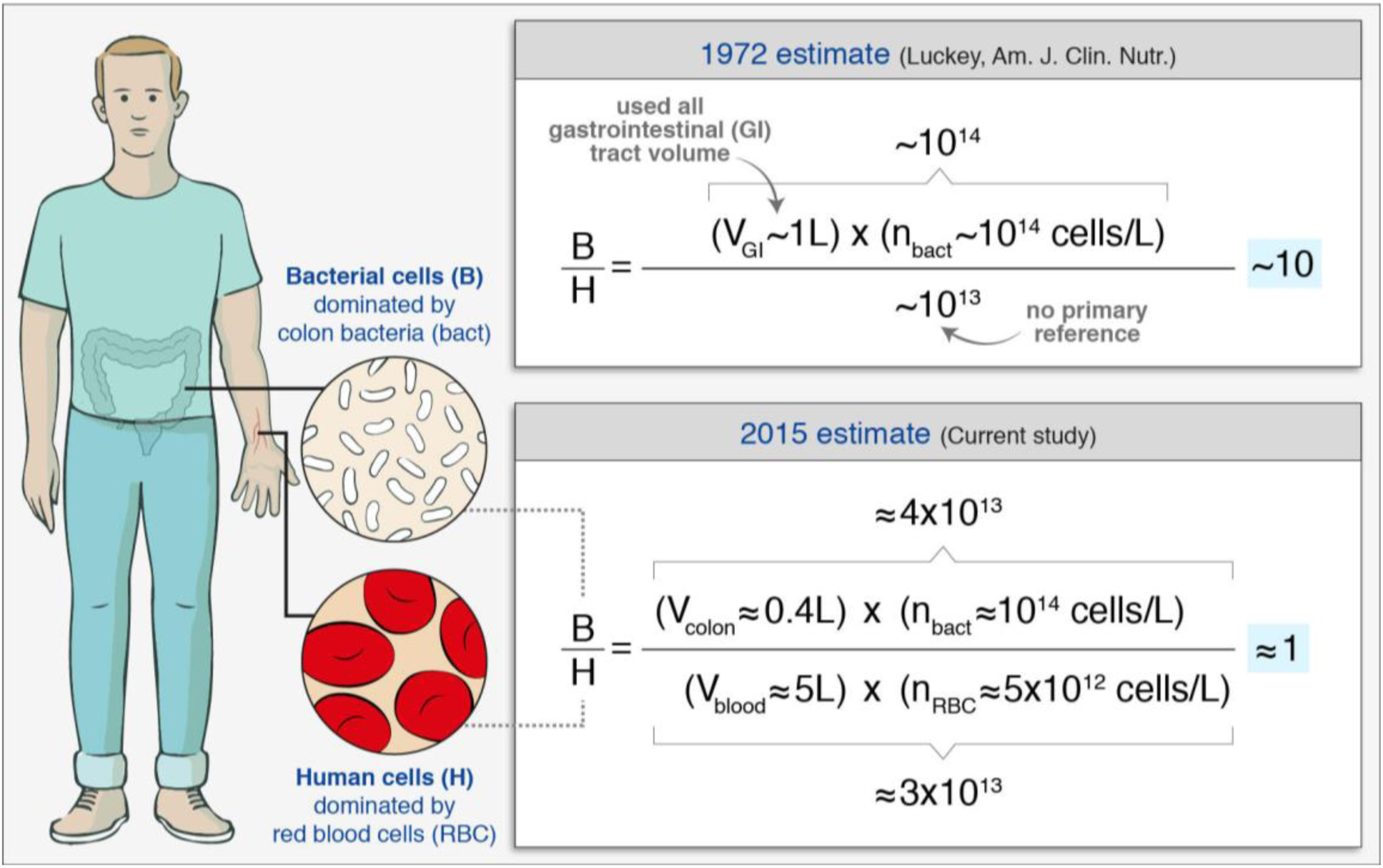
Comparison between the well cited estimate (Luckey, 1972) and the current estimate, highlighting the key four parameters identified as determining the B/H ratio in the standard man. Note that in line with formal definitions we use “~” to denote “order of magnitude” and to denote “approximately equal” (usually to better than 2-fold).

### The ratio of bacteria to human cells in the adult body

After revising both the numerator (bacteria) and denominator (human cells) in the ratio of bacteria to human cells in the body, we arrive at our updated estimate of B/H = 1.3, with an uncertainty of 25% and a variation of 53% over the population of standard 70 kg males. Comparison between the current estimate and the original estimate is illustrated in Figure 4.

We think this value and uncertainty are a much more realistic depiction that should replace the 10:1 or 100:1 values which are common in the literature at least until more accurate measurements become available.

Interestingly, if we compare the number of bacteria in the human body (3.9·10^13^) to the number of nucleated human cells (≈0.3·10^13^) we do get a ratio of about 1 to 10. We note that this ratio is the result of both the number of bacteria and the number of nucleated human cells in the body to be several times lower than in the original estimate (that did not restrict the analysis to nucleated cells).

## Discussion

Should we care about the absolute number of human cells in the body or the ratio of bacterial to human cells? A recent study shows how knowing the number of cells in different tissues can be an important indicator in understand variation in cancer risk among tissues (Tomasetti and Vogelstein, 2015). Other applications refer to the process of development and mutation accumulation. At the same time, we think that updating the ratio of bacteria to human cells from 10:1 or 100:1 to closer to 1:1 does not take away from the biological importance of the microbiota. Yet, we are convinced that a number widely stated should be based on the best available data, serving to keep the quantitative biological discourse rigorous.

The standard person used in the literature and thus analyzed above is defined as a “reference man being between 20–30 years of age, weighing 70 kg, is 170 cm in height” (Snyder et al., 1975). We now discuss the updates required in the calculation and the applicability of our conclusions to other segments in the population. To explore the effect of factors such as age, gender and body weight we focus on the four parameters (Fig. 4), which dominate any quantitatively significant deviations from the standard reference. This is because the colon bacterial count and total RBC count dominate either side of the B/H ratio. The four parameters are therefore, colon volume and bacterial density in the colon on the one hand, and hematocrit and blood volume on the other. Let us start with the gender effect. Colon volume in females is similar to that of males, 430±170 ml for a female of “standard” 1.63 m height (ICRP, 2002; Pritchard et al., 2013). As for colonic/fecal bacteria number density, there is no report in the literature of gender-specific differences. The number of red blood cells is affected by the total blood volume and by the red blood cell concentration. Red blood cell concentration is about 10% lower for females (Wakeman et al., 2007). Furthermore, blood volume is also lower by about 20–30% (Boer, 1984). Therefore, we expect the bacteria to human cell ratio to increase by about a third in females.

Proceeding to analyze infants, we note that colon bacterial density is relatively constant from infancy to adulthood (Roger and Mccartney, 2010). Colon volumes for the pediatric population, reported as 50 mL for neonates and 80 mL for one year old infants (ICRP, 2002), are derived only from comparing infant to adult daily fecal output values, and are thus less reliable and represent a knowledge gap. RBC concentration in the blood has a characteristic small temporal variation from the neonate to the elderly. RBC count values at birth are somewhat higher than for normal adults but they decrease during the first 2 months until they level at 10% lower than adult values (Zierk et al., 2015). On the other hand, the blood volume to weight of infants is 75–80 ml/kg (Howie, 2011), approximately 10% higher than normal adults. Therefore, the overall effect in terms of RBC count per body mass is smaller than 10%. In the elderly, blood volume is reduced by about 25% (Davy and Seals, 1994), while the hematocrit is essentially unchanged (Adeli et al., 2015). We therefore conclude that the effect of age on the B/H ratio is smaller than 2-fold from age one onward, and probably within the variation we estimated across the population of “standard” adult males.

Finally we analyze the effect of obesity, which is of interest in our context considering the highly intriguing links between gut microbiome and weight. Measurements of the colonic bacterial concentrations in obese individuals are similar to the ones for the reference man (Vulevic et al., 2013), indicating that the change in total bacteria number as a function of weight, is determined only by the change in colon volume. We could not find any direct measurements of the colonic volume for obese individuals in the literature yet from an indirect analysis the volume increases with weight and plateaus at about 600 ml, i.e. about 50% higher than that of the standard man value (Young et al., 2009). Moving to the number of human cells, we note that the excess body weight in high BMI individuals is mostly contributed by adipocyte hypertrophy and hyperplasia (Jo et al., 2009). Since in the reference man, fat tissue accounts for only 0.2% of the total human cell count, the added fat tissue accounts for a negligible contribution to the total human cell count. Blood volume itself increases with BMI. Because adipose tissue is not highly vascular, an increase of 100%-200% from the reference man’s body weight, to total body weights of 140–210 kg, increases the total blood volume by 40%-80% (Feldschuh and Enson, 1977). This increase in blood volume is of the same range as the increase in colonic volume in the obese and thus the B/H ratio is expected to remain within the uncertainty range we report for the “standard man”. In conclusion, the paper’s framework and general inferences on the B/H ratio are relevant for the general human population with minor quantitative differences.

We view this manuscript as a call to revitalize efforts in the direction of quantifying absolute cell content of human tissues and their commensal bacteria. Updating the ratio of bacteria to human cells from 10:1 or 100:1 to closer to 1:1 does not take away from the biological importance of the microbiota. Yet, we are convinced that a widely-stated number should be based on the best available data, serving to keep the quantitative biological discourse rigorous. Investigating whether the concentration of bacteria in stool resembles that of the colon is an important avenue along which further study is required. The analysis presented here, helps us achieve a more stable quantitative basis for discussing the cellular composition of the human body. Although we still appear to be outnumbered, we now know more reliably to what degree and can quantify our uncertainty about the ratios and absolute numbers. The B/H ratio is actually close enough to one, so that each defecation event, which excretes about 1/3 of the colonic bacterial content, may flip the ratio to favor human cells over bacteria. This anecdote serves to highlight that some variation in the ratio of bacterial to human cells occurs not only across individual humans but also over the course of the day. In addition, some medical procedures (e.g. bowel preparation before colonoscopy) decrease the bacterial colon content much more extremely than defecation and thus make the ratio significantly smaller than 1 for a period of hours to days.

We think that the kind of progression presented in this study from informative back-of-the-envelope calculations to more nuanced value estimates is of wide interest and is instructive in the quantitative training of biologists. In performing these kinds of calculations, we become intimately familiar with the limits of our current understanding and therefore, more easily highlight the best avenues for scientific progress in a particular field. What better place to start such quantitative training than by examining the contents of the human body? In doing so we can comply with the Delphic maxim of “know thyself” in a truly quantitative fashion.

## Experimental procedures

### Calculation of means, uncertainties and variation across the population

Average values were calculated using the arithmetic mean of reported values except for bacterial concentration in the colon as described below. Uncertainty in our estimates is calculated as the standard error of the mean (SEM), calculated as the standard deviation of mean values, divided by the square root of the number of reported values. For the calculation of variation across the population, the coefficient of variation (CV) was calculated as the arithmetic mean of the ratio between the standard deviation and the mean, of each of the reported values.

For the derivation of the bacterial concentration in the colon content, a large range of values was gathered from 14 different articles. The estimate for representative average value was calculated in two ways: the geometric mean and the median of the reported results. In the second case SEM was calculated from the empirical distribution using bootstrapping by the standard deviation of 1000 repeats (see SI tab BacterialConc.).

## Author contributions

R.S., S.F. and R.M. conceived and performed the study and wrote the manuscript.

## Acknowledgments

The authors are thankful for the following colleagues for helpful discussions of the manuscript: Eva Bianconi, Pascal Buenzli, Silvia Canaider, Dan Davidi, Eran Elinav, Avi Flamholz, Miki Goldenfeld, Tal Korem, Uri Moran, Nigel Orme, Rob Phillips, Silvio Pitlik, Jonathan Rosenblatt, Eran Segal, Maya Shamir, Jeff Shander, Tomer Shlomi, Rotem Sorek, Pierluigi Strippoli, Gerald Tannok, Christoph Thaiss, Jonathan Wasserman, Dave Wernick, Aryeh Wides and Lionel Wolberger.

The authors report no conflicts of interest. The authors alone are responsible for the content and writing of the paper.

This work was funded by the European Research Council (Project SYMPAC 260392); Dana and Yossie Hollander; Helmsley Charitable Foundation; Israel Ministry of Science; The Larson Charitable Foundation. R.M. is the Charles and Louise Gartner professional chair and an EMBO young investigator program member.

